# Development of an open-pollinated genetic mapping framework to facilitate the identification of QTL in apples

**DOI:** 10.1101/2025.05.08.652880

**Authors:** David Hickok, Awais Khan, Kelly Robbins

## Abstract

Genetic mapping of traits in apples (*Malus domestica*), a temperate woody perennial, is challenging due to high heterozygosity, long juvenility, and extensive spatial maintenance requirements for the populations. These factors limit the effective use of traditional quantitative trait locus (QTL) mapping methods in identifying the genetic basis of traits critical for apple production, marketability and sustainability. We explored the use of open-pollinated (OP) genetic mapping as an alternative to conventional bi-parental QTL mapping and genome wide association studies (GWAS). A QTL was simulated and the performance of a mixed linear model (MLM) and a multi-locus mixed model (MLMM) in QTL mapping accuracy was compared using a genotyping-by-sequencing (GBS) dataset of seven interspecific Royal Gala × *M. sieversii* bi-parental F1 populations. The simulation results show that the MLMM outperformed the MLM by accurately identifying the simulated QTL. Analysis of power indicated that a population size of 137 individuals is required to reach an α = 0.8 for a simulated major effect QTL. Mapping resolution analysis showed that a population size of 470-600 individuals, depending on local recombination rates, is necessary to achieve high resolution within the OP population. Simulations demonstrate the potential of OP-based genetic mapping for identifying QTL in apples, reducing the logistical challenges associated with traditional QTL mapping methods. Our results show that OP-based genetic mapping could be used to speed up the identification of novel alleles directly from diverse germplasm collections in apples.

**Core Ideas:** Bi-parental QTL mapping in apples is constrained by limited diversity, high costs, and long-term orchard space.

OP-based mapping avoids controlled pollination, clonal propagation, and large-scale F1 orchard requirements.

OP-based mapping offers an alternative to bi-parental mapping by leveraging existing germplasm and natural crosses.

OP-mapping enables QTL discovery absent full pedigree information to strategically capture broad genetic diversity.

OP F1 populations support fast QTL detection, aiding rapid breeding against emerging pathogens.

## 1 INTRODUCTION

Apples (*Malus domestica*) are temperate woody perennials grown worldwide primarily for their nutritious fruit, making them the focus of extensive breeding efforts (Khan & Korban, 2022). Apple breeding is a lengthy and challenging process, particularly when developing cultivars that combine disease resistance with desirable fruit quality (Desnoues et al., 2018; Watts et al., 2023). The long juvenile phase of apple trees, lasting between 5 to 7 years, significantly prolongs the breeding cycle of apples (Khan & Korban, 2022). Wild apple species are often the only source of disease resistance alleles, but their introduction into commercial cultivars is hindered by linkage drag, where undesirable traits, are inherited along with the desired disease resistance (Madhou et al., 2020; Stewart et al., 2023; Zhao et al., 2019). It is critical to have reliable genetic markers linked to disease resistance alleles to improve selection, reduce the linkage drag, and speed up the breeding process for developing disease resistant apple cultivars with good fruit quality (Desnoues et al., 2018; Watts et al., 2023).

In apples, bi-parental and GWAS (Genome-Wide Association Studies) genetic mapping approaches have been used to identify the genetic basis of traits and develop markers to use in apple breeding (Khan & Korban, 2022; Ma et al., 2017). Multiple molecular marker platforms including SNP arrays and GBS have been developed and utilized in QTL mapping studies (Bianco et al., 2016; Marchini & Howie, 2010; Zhang et al., 2021). While these QTL studies enabled the identification of markers closely linked to target traits, there are limitations in using these traditional QTL mapping methods in perennial, long-juvenile and highly heterozygous fruit crops. Bi-parental QTL mapping methods require, controlled pollination, maintenance of populations for several years, clonal propagation, and large orchard space, to achieve high-resolution mapping (Delgado et al., 2021; Desnoues et al., 2018; Khan & Korban, 2012; Soriano et al., 2014; Tegtmeier et al., 2023). Additionally, bi-parental mapping is constrained by the limited genetic diversity from only two parents, which restricts its utility to mapping traits within the parents and their segregating population. Whereas, GWAS leverages genetically diversity germplasm collections for high resolution mapping, but requires dense marker panels and large population sizes, making it resource-intensive for QTL mapping (Larsen et al., 2018; Ma et al., 2017; Urrestarazu et al., 2017; Watts et al., 2023; M.-Y. Zhang et al., 2021). Moreover, genotyping of GWAS panels in highly heterozygous species is often challenging, leading to ascertainment bias. This bias arises from incorrect SNP calls, low call rates for certain accessions, and missing data (Bianco et al., 2016).

OP mapping presents a promising alternative to overcome these challenges associated with traditional QTL mapping and to speed up genetic mapping in long-juvenile and heterozygous crops and forest trees (Myles et al., 2009). Genetic mapping using OP populations, where only the maternal parent is known, can take advantage of the genetic diversity and allow for the establishment of large populations with a large number of recombinants, for high-resolution QTL mapping (Kumar et al., 2016; Plomion et al., 1995). OP and multi-parent populations have been used in multiple crop species and have helped to gain insight into genetic diversity, genetic characterization, and genetic mapping. In sunflower OP populations have been used to genetically characterize various traits such as leaf morphology, head diameter and days to maturity (Moreno et al., 2013). In Pearl Millet, a collection of selfed and open pollinated individuals has been used to map grain iron and zinc content (Kumar et al., 2016) and in six-row barley multi-parent mapping has led to the detection of 23 flowering time QTL (Hemshrot et al., 2019). Beyond non woody perennial species, OP mapping has been successfully implemented in a variety of species, demonstrating its effectiveness in other heterozygous and long-lived plants.

For instance, in forest trees such as Japanese larch (*Larix kaempferi*) and maritime pine (*Pinus pinaster*), OP populations have enabled the identification of QTL associated with growth, adaptation, and disease resistance by leveraging large family sizes and extensive recombination events (Dong et al., 2024; Plomion et al., 1995).Genetic mapping based on OP populations could bridge the gap between GWAS and bi-parental QTL mapping, offering an efficient alternative for identifying marker-trait associations.

Various genetic mapping strategies have been employed for QTL mapping in both bi-parental populations and diversity collections (Khan & Korban, 2012). Interval mapping and composite interval mapping (CIM) are traditional methods used in bi-parental QTL mapping, leveraging linkage information to detect QTL positions with high power (Lander & Botstein, 1989; Zeng, 1994). These methods perform well in controlled populations but are limited by the genetic diversity of the parents. For GWAS, linear models, such as the General Linear Model (GLM) and MLM, are widely used to account for population structure and relatedness, improving the accuracy of marker-trait associations (Yu et al., 2006; Z. Zhang et al., 2010). Non-parametric approaches, such as the Kruskal-Wallis (K-W) test have also been employed for initial scans of marker-trait associations, and although it provides a simple and effective method for detecting associations, the K-W test is not robust against variance heterogeneity or imbalanced genotype categories, making it less suitable for species with complex population dynamics and diverse genetic backgrounds (Konietschke et al., 2012). To improve QTL detection and power MLMM has shown promise in increasing the resolution and accuracy of QTL mapping by incorporating multiple loci into the model while reducing false positives (Segura et al., 2012).

The effectiveness of these mapping strategies depends on several factors, including population size, marker density, and genetic variance of the traits under investigation. Larger population sizes increase the statistical power to detect QTL and improve mapping resolution (Cockram & Mackay, 2018; Y. Xu et al., 2017). Marker density is critical in capturing recombination events, particularly in species with rapid linkage disequilibrium (LD) decay, such as apples, where high-density SNP panels or GBS platforms are required (Khan & Korban, 2012; Myles et al., 2009). Additionally, the effectiveness of QTL mapping is influenced by the proportion of phenotypic variance explained by the QTL. QTLs with smaller effects require larger population sizes to achieve reliable detection, as larger populations provide increased statistical power and greater power in identifying QTL (Beavis, 1994; Hall et al., 2016). Accurately estimating the variance contributed by a QTL is crucial for understanding its impact on the trait and for genetic mapping (King & Long, 2017; S. Xu, 2003). Utilizing high-density marker platforms such as SNP arrays and genotyping-by-sequencing GBS, genetic mapping can achieve comparable QTL detection power without relying solely on traditional LM approaches (Spindel et al., 2013). Dense marker coverage enhances the resolution of association studies, enabling more precise QTL localization and marker-trait associations. This advancement allows for more cost-effective and efficient genetic mapping, particularly in highly heterozygous and complex genomes like those of woody perennials (Bianco et al., 2016).

In this study, simulated phenotypic data for an OP genetic mapping population comprised of a collection of bi-parental F1 apple populations sharing a common seed parent was used to evaluate genetic models identifying marker-trait associations. Additionally, the simulation pipeline determined the population sizes required for the genetic mapping of traits with moderate to high genetic variance explained and assessed the power and mapping resolution of the marker-trait associations across different population sizes. Our findings provide insights into optimizing OP-based genetic mappings, offering an alternative method for future genetic mapping of heterozygous crops with long juvenile phases.

## 2 MATERIALS AND METHODS

### 2.1 Mapping populations and GBS SNP dataset

A GBS dataset of seven interspecific bi-parental F1 populations was used to develop and evaluate an OP genetic mapping framework to identify QTL. Seven *M*. *sieversii* pollen donors KAZ 95 06-1 (<u>PI 600497</u>), KAZ 95 08-0 (<u>PI 692439</u>), KAZ 95 10-04F (PI 613978), KAZ 95 10-05F (<u>PI 600526</u>), KAZ 95 17-14 (<u>PI 613959</u>), KAZ 95 18-07 (<u>PI 613981</u>), and KAZ 96 05-04 (<u>PI 600601</u>) were crossed with a single maternal Royal Gala (PI651008) parent to develop this population (Figure 1). Three of these F1 populations were used and described previously in genetic mapping studies (Bai et al., 2012; Norelli et al., 2017; Tegtmeier et al., 2023). Sourcing of the paternal *M*. *sieversii* genotypes took place between 1989 and 1996 (Hokanson et al., 1997; Luby et al., 2001). These pollen parents originated from Kazakhstan and were collected and included in the *Malus* germplasm collection at the USDA-ARS Plant Genetic Resources Unit (PGRU) in Geneva, NY.

**Figure 1.**
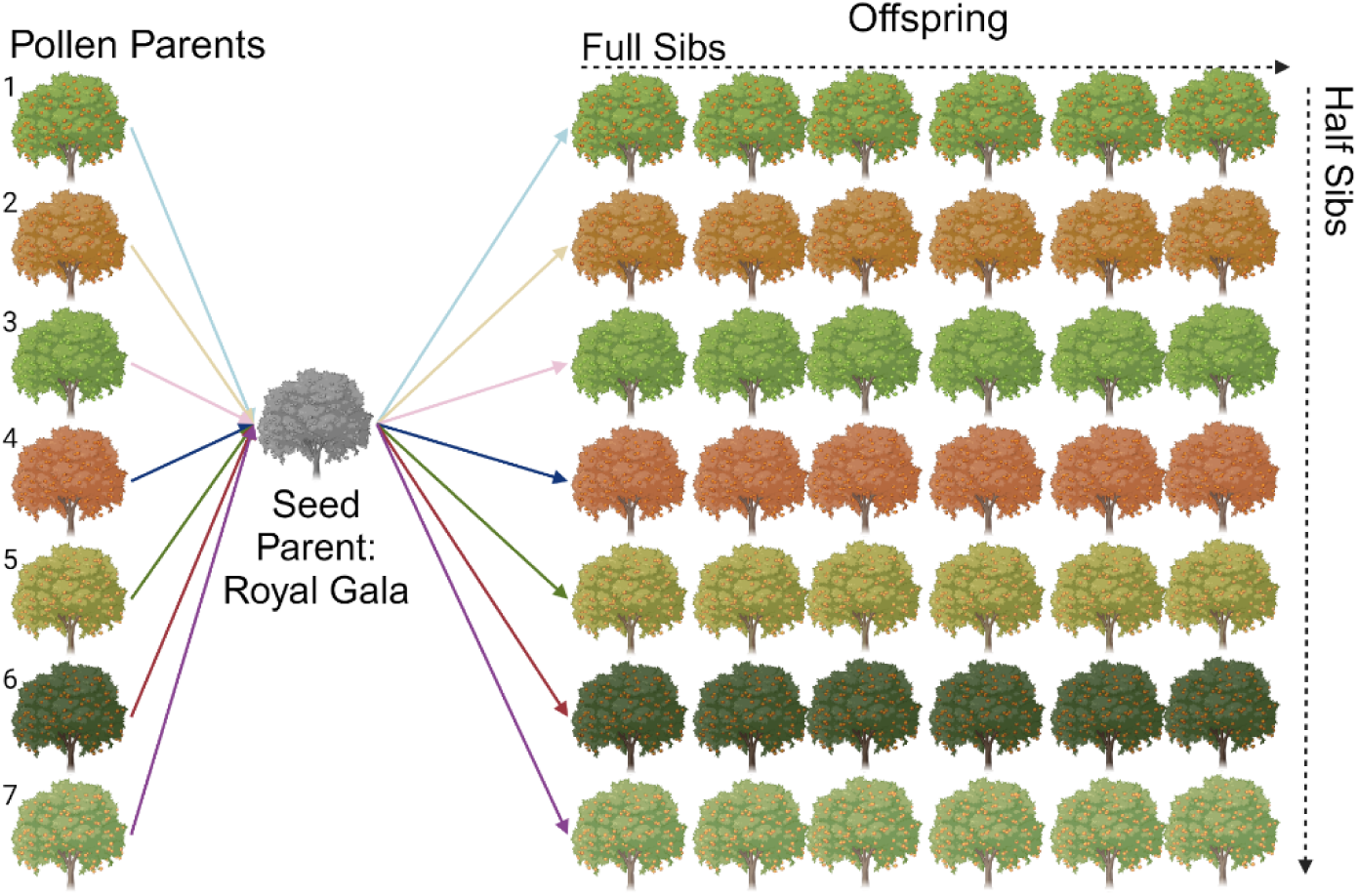
Representation of the Royal Gala (PI651008) × *M. sieversii* bi-parental F1 populations. The seven *M. sieversii* pollen parents shown on the left were crossed with the common Royal Gala seed parent shown in the middle. The arrows indicate the crosses and the resulting seven full sib populations. The F1s are aligned in rows across from the respective pollen parents with half sibs aligning in columns.

The GBS data for these populations was generated as a part of a collaborative project between USDA–ARS, Geneva, NY and Dalhousie University. The GBS genotyping protocol has been detailed in previous studies (Migicovsky et al., 2016a, 2022; Tegtmeier et al., 2023; Thapa et al., 2021). Briefly, DNA was extracted using DNeasy 96 Plant Kits (Qiagen, Valencia, CA, USA) and separate GBS libraries were prepared for each population following a modified version of the protocol developed by (Elshire et al., 2011) utilizing ApeKI, and PstI-EcoT22I restriction enzymes. Sequencing of libraries were performed on an Illumina Hi-Seq 2000 platform, with 96 samples per lane, at Cornell University (Ithaca, New York, USA). The resulting 100-bp single-end reads were filtered, and SNPs were called against the *M. domestica* GDDH13 v1.1 reference genome (Daccord et al., 2017) as described by (Migicovsky et al., 2021). The progeny are all half siblings, allowing the data to be treated as OP when analyzed in aggregate and ignoring pollen donor identity.

### 2.2 Simulation of phenotypes and QTL

Marker associations were identified using simulated phenotypes and the Royal Gala (PI651008) × *M*. *sieversii* GBS dataset. The following model representing the sampled phenotypes was used:

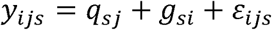

In this equation, *y_ijs_* represents the phenotypic value of the *i*^th^ plant, where *q_sj_* is the effect of QTL genotype j from simulation scenario *s*; *g_si_* is the genetic background effect of the *i*^th^ plant (polygenetic background effect for the trait of interest) sampled using the parameters for simulation scenario *s*; and *E_ijs_* is the random residual error. The QTL effect *q_sj_* was simulated to explain either 0.5 or 0.8 of the total genetic variance dependent on the simulation scenario *s*. The term *q* was simulated as 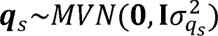 where *q*_s_ is a vector with two values representing the resistant and susceptible alleles. The genetic background effect was simulated as 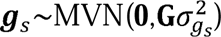, where ***g***_s_ is a vector of genetic background effects with a length equal to the sample size of simulation scenario *s*, *G* is the Genomic Relationship Matrix (GRM) calculated using the VanRaden (2008) method, and 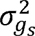 is the scalar representing the polygenic variance which would explain 0.5 or 0.2 of the total genetic variance depending on the magnitude of the genetic variance explained by the QTL effect, such that the total genetic variation remained constant across all simulation scenarios. The residual term *E_ijs_* captures unexplained variation and was sampled as 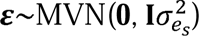, where l is the identity matrix and 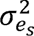 is the residual variance contributing to the phenotypic value y, with the value of 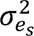 set to achieve the desired heritability of simulation scenario s. All experiments were simulated as unreplicated.

The trait was simulated such that the allelic state of the Royal Gala maternal parent was heterozygous at the locus (A/a), while all paternal *M. sieversii* parents were homozygous (a/a) at the locus. This configuration ensured the segregation of the allele unique to the seed parent within the F population, with inheritance of the QTL exclusively derived from the allelic state of the seed parent, Royal Gala. By doing so, each progeny plant inherited either the allele unique to the seed parent allele or the non-resistant allele from the seed parent, allowing for segregation between resistant and non-resistant phenotypes based on the presence or absence of the seed parent allele.

#### 2.2.1 Simulation parameters

To evaluate the effect of population size on QTL detection a multi-step procedure was used. First, the GBS dataset was imputed using BEAGLE (Browning et al., 2018) and the bi-parental GBS dataset was used to calculate a GRM for all individuals. Next, the MLMM was run 15 times, each time on a reduced population size. For each of these population sizes, phenotypes were simulated under the predefined QTL parameters (Table 1). The simulated phenotypes were then analyzed with the MLMM to obtain marker-trait associations. We repeated this process 40 times for each reduced population size, generating new simulated phenotypes in each repetition. Finally, the identified marker-trait associations were compared to the location of the simulated QTL to calculate both power, false discovery rate (FDR), and fine-mapping resolution (Figure 2).

**Figure 2.**
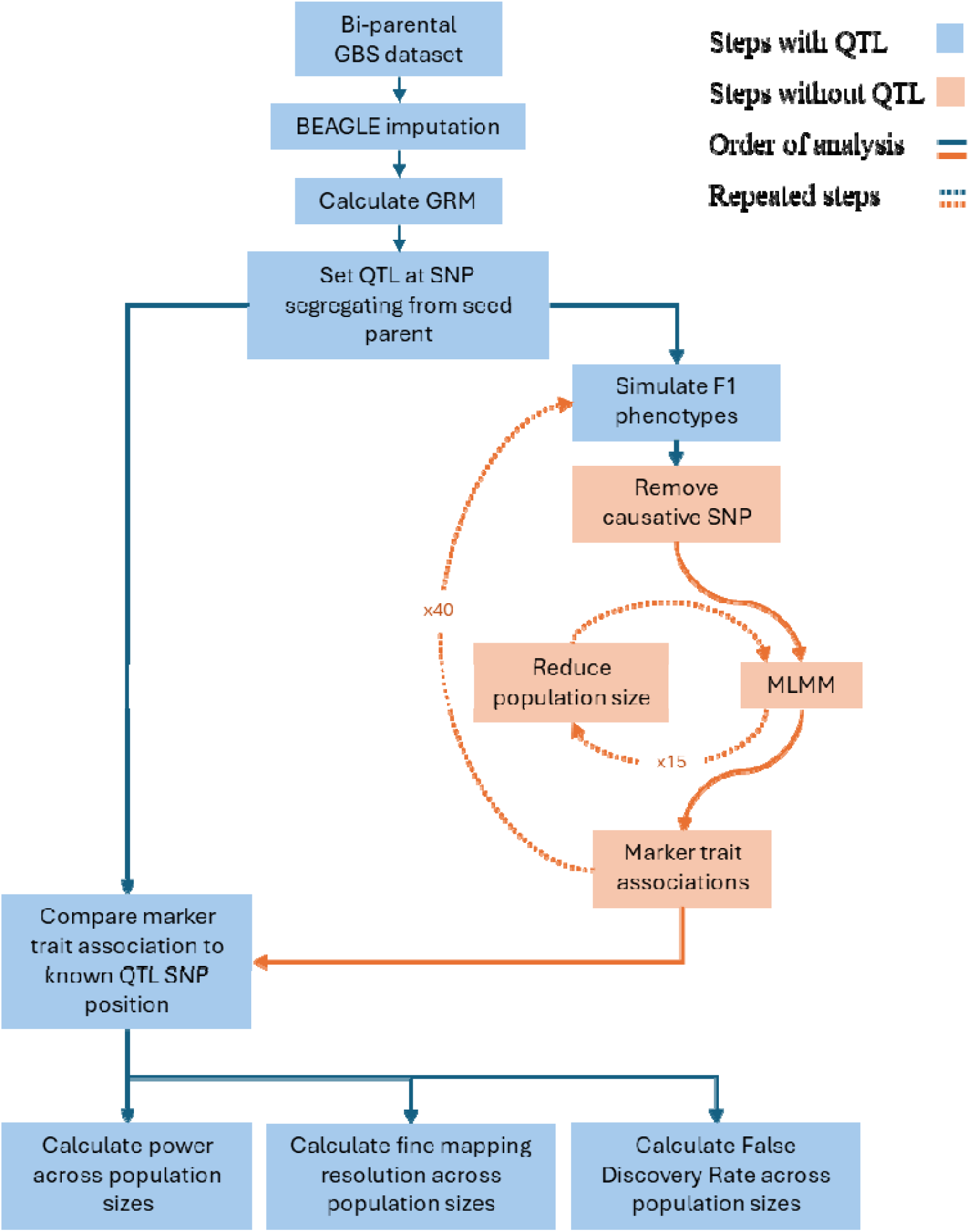
Workflow used to calculate power, FDR, and determine resolution of marker-trait associations. Blue arrows indicate the sequence of steps involved in simulating phenotypes and analysis of marker-trait associations including the QTL. Orange arrows indicate the direction of simulation and analysis that where the QTL is masked and do not include the simulated QTL. Dashed lines indicate steps that are repeated. Orange boxes indicate steps excluding the QTL. Blue boxes indicate steps that include the QTL. First, the bi-parental GBS dataset is used to calculate a genomic relationship matrix (GRM). Next, the MLMM is run 15 times with a reduced population size each run. Phenotypes are simulated 40 times for the population. The process of obtaining marker-trait associations from these simulated phenotypes is repeated 40 times, with new phenotypes being simulated for each repetition. Marker associations are compared to the position of the simulated QTL to calculate power, FDR, and fine mapping resolution.

**Table 1.** Simulation parameters for all scenarios for model comparison and power, false discovery rate (FDR), and resolution comparisons. Scenarios 1 and 2 were located on LG 3 with a heritability of 0.8, and genetic variance of 0.8, while Scenario 6 was located on LG 7 with a heritability of 0.89 and genetic variance of 0.5, based on parameters for a previously identified fire blight resistance QTL, *MSV_FB7*. For scenarios 4 – 7, the heritability was set to 0.5, with corresponding genetic variances of 0.5 or 0.8, respectively. Simulations were conducted on two linkage groups (LG 7 and LG 1) categorized with higher (LD = 0.3) or lower (LD = 0.03).

| Simulation Parameters: MLM and MLMM |  |  |  |  |  |  |
| --- | --- | --- | --- | --- | --- | --- |
| Scenario | Heritability | Genetic Variance | Linkage Group | SNP |  | Model |
| 1 | 0.80 | 0.80 | LG 3 | SLG3_PHT_2552347 |  | MLM |
| 2 | 0.80 | 0.80 | LG 3 | SLG3_PHT_2552347 |  | MLMM |
| 3 | 0.89 | 0.50 | LG 7 | SLG7_PHT_27422647 |  | MLMM |
| Simulation Parameters: Power, FDR, Resolution |  |  |  |  |  |  |
| Scenario | Heritability | Genetic Variance | Linkage Group | LD Level | SNP | Model |
| 4 | 0.50 | 0.50 | LG 7 | 0.30 | SLG7_PHT_27636570 | MLMM |
| 5 | 0.50 | 0.80 | LG 7 | 0.30 | SLG7_PHT_27636570 | MLMM |
| 6 | 0.50 | 0.50 | LG 1 | 0.03 | SLG1_PHT_18796150 | MLMM |
| 7 | 0.50 | 0.80 | LG 1 | 0.03 | SLG1_PHT_18796150 | MLMM |

To assess how the proportion of genetic variance explained by a QTL marker-trait associations, we simulated phenotypes from the GBS dataset using two values of *V_G_* (0.5 and 0.8). In addition, we incorporated a heritability of 0.5 to represent scenarios where disease resistance traits in apple can exhibit moderate heritability, as reported for fire blight (Kostick et al., 2021), apple scab (Bus et al., 2002), and powdery mildew (Sestras et al., 2011). Accordingly, the simulated QTL in simulation scenarios (4-7) approximate the moderate-heritability levels that have been reported for some disease resistance traits in apple (Table 1).

### 2.3 Imputation of genotypic data

To ensure the correctness of genotype calls in the F populations, we first identified and removed impossible allelic states based on parental genotypes. Specifically, the maternal parent (*Royal Gala*) was heterozygous at the locus of interest, while all paternal parents (*M. sieversii*) were homozygous recessive and did not carry the desired allele. Consequently, any progeny genotype that did not conform to these parental configurations (e.g., A/A when the paternal parent is a/a) was deemed impossible and set to missing ‘NA’.

To address the resulting missing data, we employed BEAGLE for genotype imputation. BEAGLE was utilized without incorporating parental genotype information, treating the dataset as if parentage were unknown. This approach leveraged the LD patterns and haplotype structures within the population to accurately infer the missing genotypes agnostic to potential parentage. The filtered GBS dataset was formatted according to BEAGLE’s requirements and imputed using standard parameter settings, including default iteration counts and effective population size estimates.

### 2.4 QTL localization

To investigate the ability to fine map QTL in OP populations of apples, QTL were simulated within genomic regions defined by differing levels of LD. Two (25kb) regions, one with low average LD (0.03) across the region and one with higher LD (0.30) across the region were chosen, representing higher and lower recombination regions. The workflow of the simulation is represented in (Figure 2).

### 2.5 Model selection for genetic mapping

Evaluation of different mapping approaches was performed to determine the effectiveness of two different mixed linear models: the MLM and the MLMM. The MLM accounts for population structure by incorporating a kinship matrix, thereby controlling for relatedness among individuals. In contrast, the MLMM extends the MLM by additionally accounting for confounding background genetic effects through the inclusion of multiple genetic markers as fixed covariates using a stepwise regression approach (Segura et al., 2012). Both models were implemented using the Genome Association and Prediction Integrated Tool (GAPIT v3.0) within the R (version 4.2.2) environment (Lipka et al., 2012). All genotypic and simulated phenotypic data were formatted according to GAPIT standards. Comparison of MLM and MLMM for a single QTL of heritability of 0.8 and percent genotypic variation explained of 0.8, similar to that of MR5, a fire blight resistance gene located on linkage group (LG) 3 (Peil et al., 2007). Additionally, the simulation parameters were informed by findings from previous research on the genetic architecture of the bi-parental subpopulation M4591, specifically focusing on *MSV_FB7*. *MSV_FB7* is a fire blight resistance locus with an effect size of 0.5 and a heritability of 0.89, located on LG 7 (Tegtmeier et al., 2023).

### 2.6 Linkage mapping and OP mapping comparison

The heritability, variation explained, and location of the QTL was set similar to the *MSV_FB7* fire blight resistance QTL (Tegtmeier et al., 2023) for simulations and to compare an OP-based genetic mapping method to a bi-parental QTL mapping method. Heritability was set to 0.89 and variance explained by the locus was set to 0.5. Evaluation of different mapping approaches was performed to determine the effectiveness of the MLM and the MLMM. Based on initial comparisons between the MLM and MLMM (Scenarios 1–3), we selected the MLMM as our primary mapping approach for subsequent simulations. In Scenarios 1 and 2, we modeled a major-effect QTL (*V_G_* = 0.8 and h^2^ = 0.8) on LG 3, reflecting parameters similar to *MR5*, a known fire blight resistance locus (Peil et al., 2007). Scenario 6 further examined a moderate-effect QTL (*V_G_* = 0.5 and h^2^ = 0.89) on LG 7, reflecting parameters similar to *MSV_FB7*, another known fire blight resistance locus (Tegtmeier et al., 2023). Having identified the MLMM as superior in these reference cases, we next evaluated how this method performs over multiple simulated scenarios 4 - 7 where the heritability was set to 0.5 and genetic variance was set to either 0.5 or 0.8 (Table 1). These lower heritabilities and varying LD levels (LG 1 vs. LG 7) provide a broader assessment of OP-based genetic mapping in apple, as moderate-effect loci are common in breeding programs and LD landscape will be different depending on the genetic architecture surrounding the QTL.

### 2.7 Effect of population size on the OP-based genetic mapping

Individuals were filtered out of the dataset to determine the effect of population size on the OP-based genetic mapping framework by calculating the power of correctly identifying significant SNPs using the MLMM. The population was filtered such that the proportion of individuals from each bi-parental population remained constant. This was done by removing an equal percentage of individuals from each of the bi-parental populations before running the MLMM with the simulated phenotypes. To maintain an equal proportion of each of the seven bi-parental populations within each simulation a percent of each population was excluded from the dataset for every round of simulations as follows *N_i_* = [*N_total_* - *N_total_* X *p_i_*] where *N_i_* is the resulting population size, *N_total_* is the total population size, and *p_i_* in {0.35,0.40,0.45, …, 0.95}.

### 2.8 Power analysis and mapping resolution

Statistical power was evaluated to determine the probability of correctly identifying a true QTL under the conditions of the simulations. Power was calculated for each population size as the proportion of simulation iterations in which at least one SNP within a 2Mb range of the true causative QTL was identified as significant. The power analysis involved simulating phenotypes with known effects and performing marker-trait associations using the MLMM. SNPs were considered significant if their *p*-values fell below the Bonferroni-corrected threshold of 0.05. Power was then calculated as the ratio of simulation runs where the QTL was detected to the total number of simulation runs. The equation used is given below:

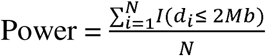

Where *N* is the total number (40) of simulations for each population size, *d_i_* is the distance between the closest significant SNP identified in iteration *i* and the true causative QTL. Where *I*(*d_i_* < 2*Mb*) is an indicator function that equals 1 if at least one SNP within a 2Mb range of the simulated QTL was identified as significant in iteration *i*, and 0 if there was no SNP within the 2Mb range of the simulated QTL. This analysis was conducted across different scenarios that varied in genetic variance, LD levels, and population sizes. By evaluating power under these varying conditions, we were able to determine the effectiveness of our OP-based genetic mapping framework in reliably identifying true QTLs. The resulting power estimates provided a comprehensive understanding of the framework’s capability to detect genetic associations under diverse genetic architectures and population structures. Mapping resolution was calculated as the distance to the causative SNP:

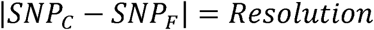

where *SNP_C_* is the causative SNP and *SNP_F_* is the furthest identified linked SNP in base pairs. For each simulation the distance between SNP_C_ and SNP_N_ was calculated and the mean distance across all iterations and was calculated to determine the average mapping resolution.

### 2.9 False discovery rate

The FDR was calculated to determine the proportion of false positive SNPs among those identified as significantly associated with the simulated QTL. The FDR was determined using the formula:

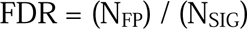

where N_FP_ represents the number of false positive SNPs, those identified as significant but located outside a 2Mb window surrounding the simulated QTL and N_SIG_ denotes the total number of SNPs identified as significant.

## 3 RESULTS

### 3.1 Imputation

Across the 915 F individuals and 34,280 genotyped sites, we identified a total of 2,603,916 allelic states (8.3% of all calls) that were biologically impossible given the parental configurations described above. These calls were subsequently set to missing ‘NA’, which combined with the existing 1,927,671 missing sites, yielded 4,531,587 missing data points in total. This quantity accounted for approximately 14.45% of the 31,361,200 genotype calls in the dataset.

### 3.2 Model comparison between MLM and MLMM

A simulated QTL with parameters identical to the previously reported fire blight resistance QTL, *MSV_FB7* i.e h^2^ = 0.89, *V_G_* =0.5, which segregates in the bi-parental population GMAL 4591 was successfully identified using the MLMM on linkage group 7 (Figure 3c). The associated SNP, SLG7_PHT_27422647, showed a highly significant association (*p*-value = 7.95e-146) with the trait and this SNP falls within the previously identified QTL region for *MSV_FB7*, spanning from markers SLG7_PHT_22699693 to SLG7_PHT_27725770. The comparison between the MLM and the MLMM for a single QTL was done using simulation scenarios 1 – 3 (Table 1) highlighting distinct patterns in the association profiles of each model. The MLM identified nine SNP marker peaks across nine different linkage groups, suggesting erroneous associations with the simulated QTL. In contrast, the MLMM isolated a single, strong association precisely at the simulated QTL on LG 3, reducing the unrelated associations found with the MLM. Figure (3a) demonstrates that the MLM’s associations were dispersed across multiple LGs, potentially due to population structure or unaccounted background genetics, whereas (Figure 3b) shows the MLMM pinpointing only the intended QTL region. These results illustrate that MLMM approach provides more targeted associations by controlling for confounding effects. Consequently, the MLMM proved more effective in minimizing spurious associations, enhancing the detection of the simulated QTL. This reduction in extraneous peaks validated MLMM’s suitability for the genetic structure of open-pollinated apple populations and led to its selection for all subsequent simulations.

**Figure 3.**
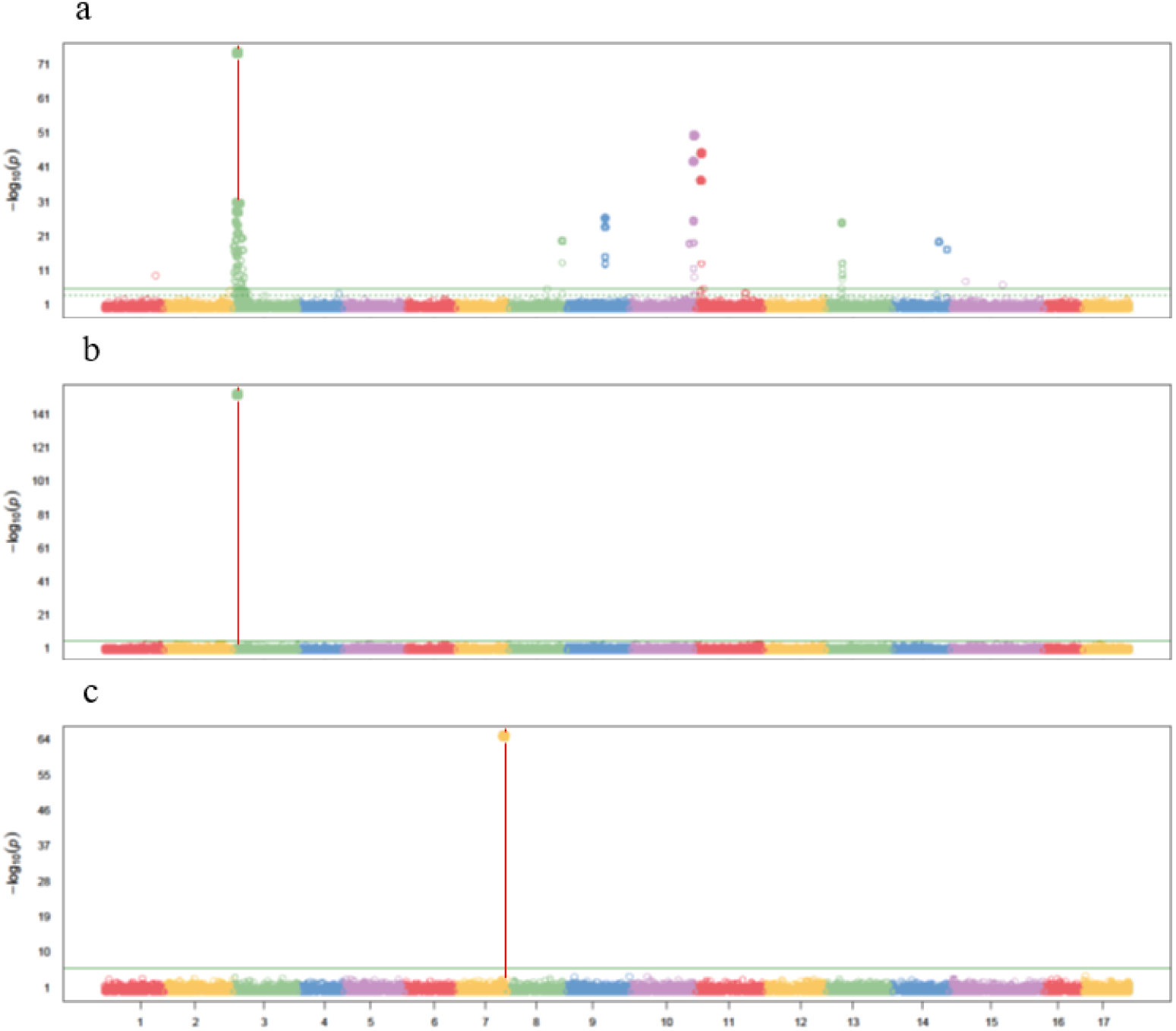
(a, **b)** Manhattan plots comparing marker-trait associations using two different models, mixed linear model (MLM) and multi locus mixed model (MLMM) for simulation scenario 1 in panel a simulation scenario 2 in panel b using the dataset from the Royal Gala (PI651008) × *M. sieversii* bi-parental F1 populations. The dashed line is the genome wide significance threshold, and the solid line is the Bonferroni corrected threshold of 0.05. X-axis shows the chromosome number, and the y-axis is −log10(p) value. SNPs are represented by different colored points with the colors indicating the linkage group (LG) the variant is located. A red vertical line indicate the location of the simulated QTL. is the heritability and is the genetic variance of the simulated QTL. The SNPs detected by the MLM were on 9 different LGs and the SNP detected by the MLMM was the same as was selected for the simulated QTL SLG3_PHT_2552347. **(c)** Manhattan plot of marker – trait association results for simulation scenario 3 with the same h^2^, *V_G_*, and location as a previously identified fire blight resistance QTL on LG07. The Manhattan plot for the simulated QTL shows the linked SNP SLG7 PHT 27422647 within the previously identified *MSV_FB7* QTL region (Tegtmeier et al. 2023) between markers SLG7_PHT_22699693 to SLG7_PHT_27725770.

### 3.3 Power analysis, false discovery rate, and fine mapping resolution

### 3.4 Statistical power under different levels of *V_G_*

Over different proportions of genetic variations explained by the simulated QTL (*V_G_* =0.5 and *V_G_* =0.8) on both LG 1 and LG 7, it was observed that the number of individuals in the population required to achieve a statistical power of 0.8 decreased with a higher *V_G_* and lower recombination rate. For the simulated QTL on LG 1, the major effect QTL with *V_G_* =0.8 required a population size of 183 individuals to reach the beta threshold 0.8 (Figure 5). In contrast, the moderate effect QTL on LG 1 with *V_G_* =0.5 required a larger population size of 320 individuals to achieve the same alpha threshold. For the simulated QTL on LG 7, the major effect QTL with *V_G_* =0.8 required a population size of 137 individuals to reach a power of 0.8, while the moderate effect QTL required a population size of 274 individuals to achieve the same power (Figure 5, Table S3).

The power analysis showed that the ability to detect simulated QTLs varies with genetic variance explained, chromosomal location, and population size. Notably, the simulated QTL on LG 7 required smaller populations than the simulated QTL on LG 1 to reach the same power, suggesting that LD influences marker trait associations. Published linkage maps of apple have shown that recombination rates can vary significantly among linkage groups, thus impacting the extent of LD decay (Di Pierro et al., 2016). For major effect QTLs, *V_G_* =0.8, the statistical power reached the threshold of 0.8 at smaller population sizes compared to moderate effect QTLs, *V_G_* =0.5.

For both LG 1 and LG 7 the marker-trait associations were found to be closer to the causative SNP when the distance to the causative SNP is averaged across the 40 repeated simulations, particularly for greater population sizes and the proportion of genetic variation explained by the QTL. At the population size for which each simulation scenario reached the power threshold, the resolution to the causative SNP differed. For the simulated QTL on LG 1 with *V_G_* 0.8, the average distance to the causative SNP was 1.29Mb and 1.06Mb for the simulated QTL on LG 7. For the simulated QTL on LG 7, the major effect QTL required a population size of approximately 183 individuals to achieve a resolution within 1Mb of the causative marker. The moderate effect QTL on LG 7 would require a population size of 366 individuals to reach a similar level of resolution (Figure 4, Table S1). The FDR of the simulated QTL varies according to population size, linkage group (LG 1 vs. LG 7), and QTL effect size (moderate vs. major) (Figure 6, Table S2). We conducted an FDR analysis to measure the proportion of false positives among the significant SNPs in each scenario (Table 1). Overall, larger sample sizes reduce the FDR across all linkage groups and effect sizes. For instance, scenario 6 (LG 1 moderate) remains above 0.1 at smaller populations but drops below FDR = 0.1 around n = 229 (FDR ≈ 0.098). By contrast, Scenario 7 dips under 0.1 by n = 183 (FDR ≈ 0.047) yet exhibits some variability around that population size (e.g., FDR ≈ 0.17 at n = 183 and decreasing to 0.048 by n = 229). Additionally, Scenario 5 crosses the 0.1 threshold at around n = 183 (FDR ≈ 0.024), underscoring its quicker drop in false discoveries compared to the LG 1 moderate scenario. These results confirm that major-effect QTLs (*V_G_* = 0.8) can meet an acceptable FDR threshold at smaller population sizes, whereas moderate-effect QTLs (*V_G_* = 0.5) require more individuals to achieve the same level of stringency.

**Figure 4.**
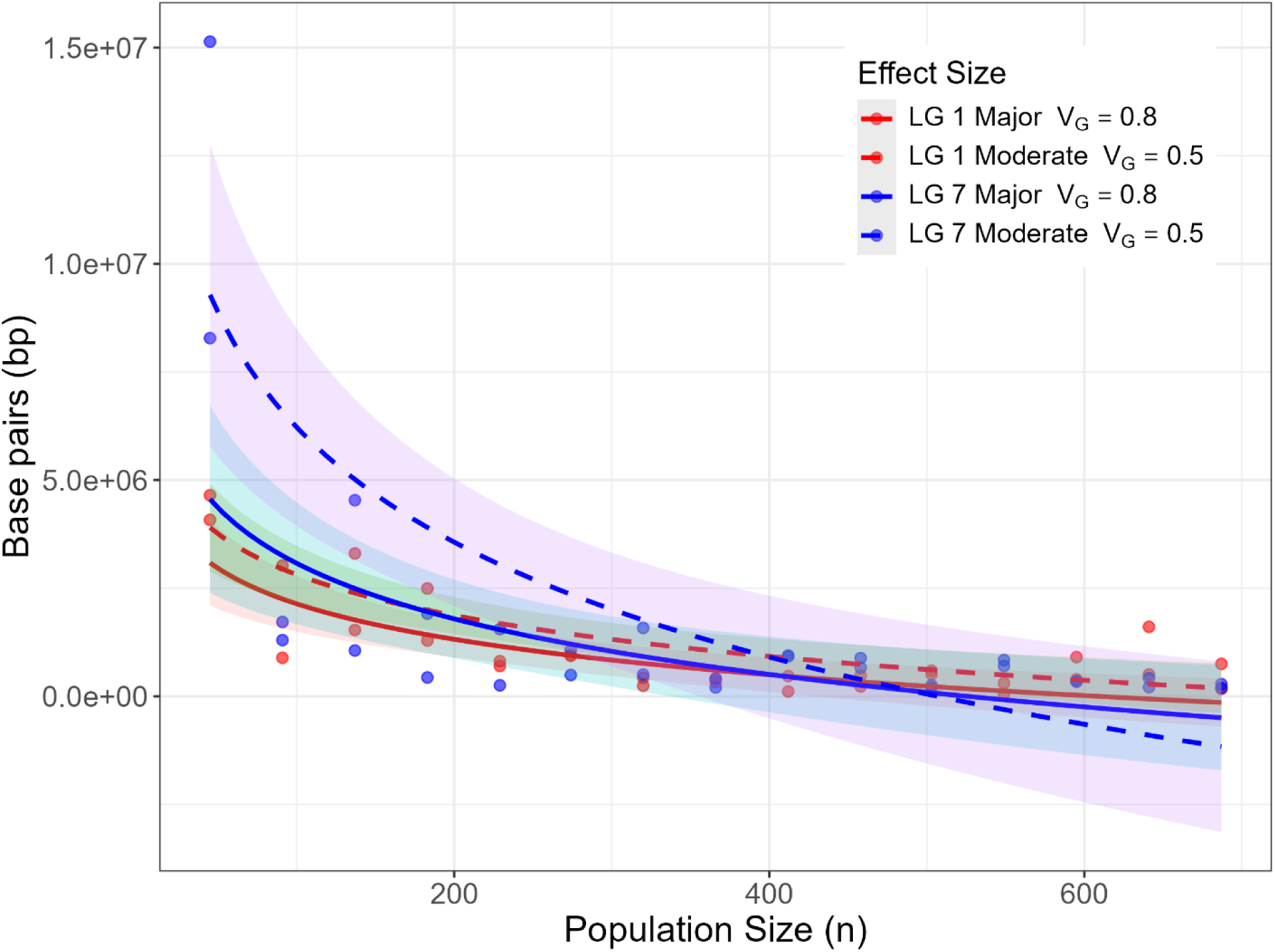
Resolution as a measurement of average distance (y-axis) between significant SNPs and the simulated QTL using an MLMM. The QTL was simulated on two linkage groups (LG 1 and LG 7) under two levels of genetic variance (0.5 = “moderate” and 0.8 = “major”) across 15 population sizes x-axis. Each point represents the average distance between the furthest significant SNP and the causal SNP over 40 simulation replicates at a given population size (x-axis). Lower values indicate greater mapping precision. Log-linear trend lines with 95% confidence intervals (shaded ribbons) illustrate overall resolution gains with increasing sample size. Scenario 6 is indicated by LG1 Moderate (red, dashed), Scenario 7 by LG1 Major (red, solid), Scenario 5 by LG 7 Moderate (blue, dashed), and Scenario 7 by LG 7 Major (blue, solid). Under LG 1 Major (Scenario 7), resolution improves from 4.65Mb at n = 45 to 175kb at n = 687, while LG7 Major (Scenario 5) improves from 8.28Mb at n = 45 to 280kb at n = 503. LG 7 Moderate (Scenario 4) improves from 15.1Mb at n = 45 to 192kb at n = 687 and LG 1 Moderate (Scenario 6) improves from 4.1Mb at n = 45 to 749kb at n = 687. Overall, resolution improves more rapidly with increasing population size across all scenarios.

**Figure 5.**
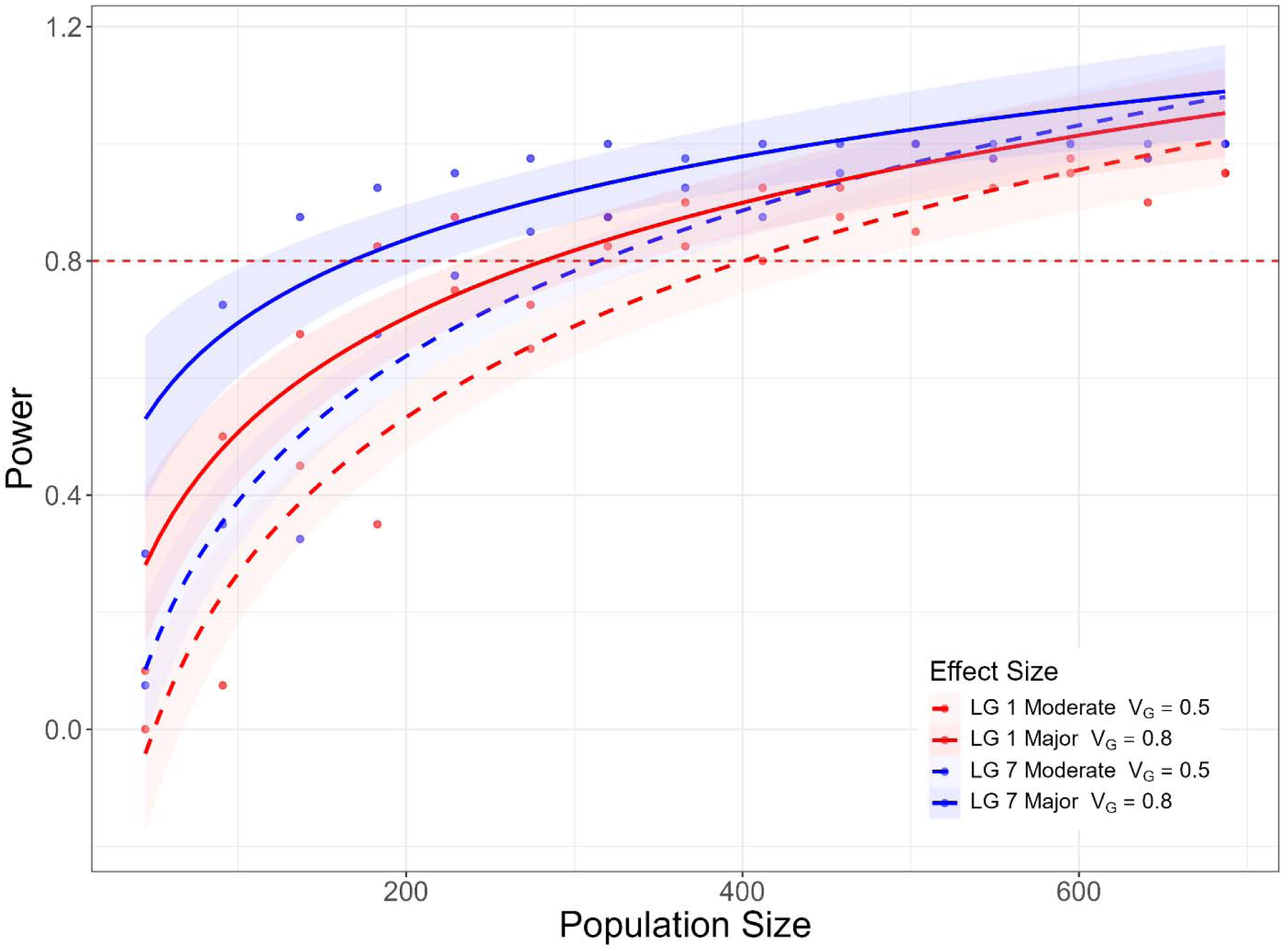
Analysis for major and moderate effect QTL simulated on either LG 1 or LG 7. The x-axis of the graph is the population size and the y-axis is the power of the multi locus mixed model (MLMM) marker - trait associations across 40 simulations for each of the 15 population sizes. The power of the marker-trait associations for the QTL simulated on LG 1 are colored in red, while the power of the marker-trait associations for the QTL simulated on LG 7 are colored in blue, moderate effect QTL (*V_G_* = 0.5) have dashed lines (with shaded confidence intervals), and major effect QTL (*V_G_* = 0.8) have solid lines (with shaded confidence intervals). The QTL was simulated with two genotypic variances explained: 0.8 and 0.5 and a phenotypic heritability of 0.5. A dashed red line indicates an alpha value of 0.8. The marker trait associations for (Scenarios 5 and 7) the QTL with *V_G_* = 0.8 reached the alpha threshold of 0.8 at a population of 183 for LG 01 (Scenario 7) and for a population of 137 for LG 7 (Scenario 5). The marker trait associations for (Scenarios 4 and 6) the QTL with *V_G_* = 0.5 reached the alpha value of 0.8 power at a population of 320 for LG 1 (Scenario 7) and a population of 274 for LG 07 (Scenario 6).

**Figure 6.**
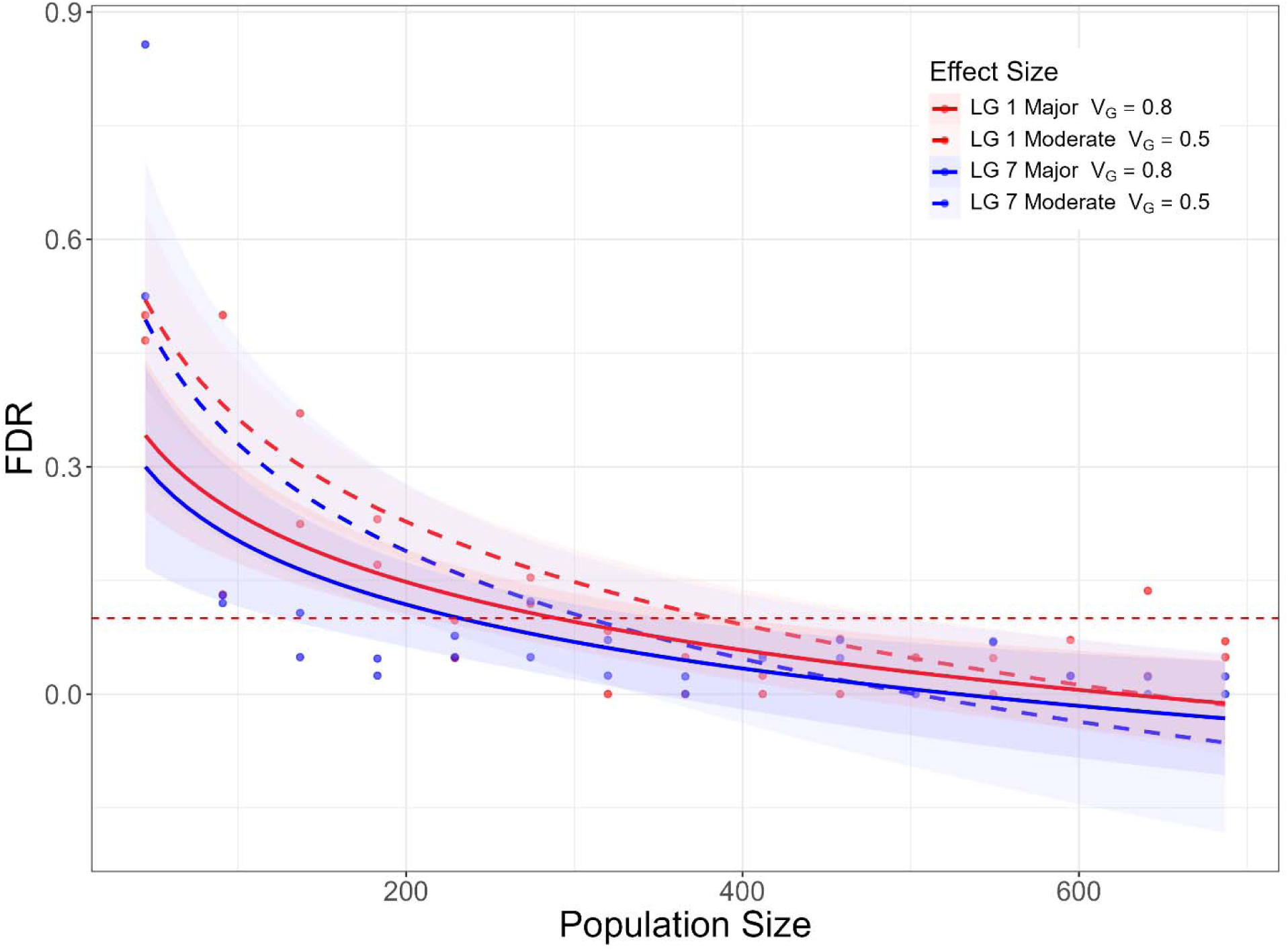
False discovery rate (FDR) displayed on the y-axis for identifying marker-trait associations of a simulated QTL using a MLMM. The QTL was simulated on two linkage groups (LG 1 and LG 7) under two levels of genetic variance (0.5 = “moderate” and 0.8 = “major”) over 15 population sizes. Each point represents the average FDR at a given population size over 40 iterations of simulating the QTL. The dashed reference line at FDR = 0.1 highlights the threshold for acceptable false discovery using the MLMM. Scenario 7 is indicated by LG 1 Major (red, solid), Scenario 6 is indicated by LG 1 Moderate (red, dashed), Scenario 5 is indicated by LG 7 Major (blue, solid), and Scenario 4 is indicated by LG 7 Moderate (blue, dashed). Scenario 7 drops below FDR = 0.1 at n = 229, while Scenario 6 falls below 0.1 at n = 329. Scenario 5 drops below FDR = 0.1 at n = 137 and Scenario 4 crosses FDR = 0.1 at n = 183, reflecting the simulated QTL on LG 7 falls under the FDR threshold of 0.1 at smaller population sizes. Additionally, larger sample sizes systematically reduce FDR across all linkage groups and effect sizes.

## 4 DISCUSSION

We developed an OP based genetic mapping framework and successfully identified simulated QTL using the genetic variation present in a half-sib F1 apple population. The results demonstrate the MLMM outperformed a standard MLM in detecting the simulated QTL. Using the MLM the simulated QTL on LG 3 was identified, but several false positives were observed across eight LGs: 1, 8, 9, 10, 11, 13, 14 and 15. MLM can yield spurious associations when background genetic effects are insufficiently accounted for, resulting in false positives that obscure the true marker-trait relationships. This has been seen in structured populations of *A. thaliana*, apples, and humans (Atwell et al., 2010; Migicovsky et al., 2016b; Platt et al., 2010). The marker-trait associations using the MLMM performed better than the MLM due to its ability to better account for multiple associated markers by including multiple loci through a stepwise approach (Segura et al., 2012). Populations with complex genetic structures can lead to spurious marker-trait associations (Newman et al., 2001), like in OP apple populations. While MLMs are effective at correcting for population structure by modeling it as a random effect (Kang et al., 2010; Yu et al., 2006). MLM test one marker at a time and can miss the combined effects of multiple loci with small individual contributions. The MLMM addresses this limitation by allowing for the joint identification of multiple loci influencing the trait, enhancing the power to detect maker-trait associations while combining the advantages of MLM in correcting for population structure with the ability to model multiple effects jointly (Segura et al., 2012). Therefore, MLMM provided a more accurate mapping of the simulated QTL in our study and identified a significant marker-trait association in all scenarios (2 – 7) utilizing the MLMM. The GBS dataset presented challenges due to improper variant calls in some of the F1 progeny in the populations. Approximately 8% of the sites had allelic states that weren’t possible in the bi-parental population when individuals’ allelic states were considered with respect to their parentage. BEAGLE allowed to impute agnostically to the parentage, however, the determination of impossible allelic states of some progeny could only be done because the parentage of the seven bi-parental populations were known. In an OP population that has unknown pollen donors, the ability to discover incorrect allelic states within the resulting mapping population, would require additional analysis, potentially employing the use of PCA or haplotype tracking (Hofmeister et al., 2022). Sequencing depth in different genotyping platforms poses another potential issue for genetic mapping in OP populations. Studies utilizing low depth sequencing data require careful consideration of structure, relatedness, and parentage for accurate imputation (Chan et al., 2016; Wang et al., 2024). A framework to reduce the resources needed would require efficient methods for accurate imputation if lower depth sequencing were to be used in F1 OP genetic mapping populations. Our results also show the importance of balancing population size and LD landscape to accurately map QTL with a range of effect sizes. Genetic mapping of a QTL on LG 1 with an average LD of 0.03 in a 25kb region around the QTL showed a convergence with the power threshold at a population size of 183 F1s compared to the moderate effect QTL requiring a population size of 320 F1 individuals to converge with the power threshold. Mapping a QTL on LG 7 with an LD block with an average LD of 0.3 in a 2kb region around the QTL showed a convergence with the power threshold at population size of 137 individuals. The moderate effect QTL required a population size of 274 individuals to converge with the power threshold. The difference in population sizes required to precisely map the QTL within the population could be due to the differing levels of genetic variation explained by the QTL, where lower effect QTL require greater mapping populations than larger effect QTL to accurately detect (Ball, 2005). This is due to the fact that QTL explaining less variation in the phenotype will have phenotypic variation from other sources, like environmental conditions and background genetic effects. In contrast, QTL that explain more genetic variance can be detected with smaller mapping populations, as the basis for phenotypic variation is due to the QTL, and much less by environmental and other polygenetic effects generating a stronger signal in marker-trait associations. The difference in population sizes required to identify marker-trait associations in contrasting strengths of LD blocks is likely due to the degree of linkage between the QTL and markers. LD and mapping population size were also important in the resolution of marker-trait associations with the simulated QTL. Over the 40 repeated QTL simulations across population sizes, the distance to the QTL decreased at a faster rate with increasing population sizes for the QTL simulated in a higher LD region, however, the distance to the QTL was closer when the QTL was in a lower LD region. Differences in fine-mapping capacity stem not only from LD strength but also from varying local recombination frequencies across different chromosomes, which can create genomic hot spots for recombination and thus unevenly affect QTL resolution (Myers et al., 2005). For a major effect QTL on LG 1 and the population size of 687 the distance to the QTL was 175kb, and the distance to the major effect QTL on LG 7, which was in a region of higher LD was 280kb. For the minor effect QTL fine mapping resolution on LG 1 was 749kb and 192kb on LG 7 for the population size of 687. The resolution of the marker – trait associations for the moderate effect QTL were similar, in that the lower LD region could map closer to the QTL than the higher LD region. Markers in higher LD with the QTL have greater marker-trait associations (Flint-Garcia et al., 2003). However, QTL with tightly linked markers are more difficult to fine map and require greater population sizes to distinguish between markers tightly linked with the QTL (Y. Xu et al., 2017). The analysis of the FDR for the simulations underscores the need to balance sample size with QTL effect size to control the proportion of false positives (Figure 6). Controlling FDR is essential for reliable marker-trait association studies (Storey & Tibshirani, 2003). We found that moderate-effect QTLs exhibit higher FDRs at smaller population sizes, whereas major-effect QTLs can achieve acceptable FDR thresholds more quickly. This pattern emphasizes that both effect magnitude and population size are important factors to consider for minimizing spurious associations, complementing the improvements in accuracy already provided by MLMM. Paternal parent inference remains an issue with the proposed method of OP based genetic mapping, particularly when using sequence-based genotyping. A two-stage approach focusing on robust genotyping of potential pollen-donors and imputing based off a subset, likely to be contributing to the OP F1 population could improve accuracy despite having unknown paternal lines. In outcrossed, highly heterozygous species like apple, low sequencing depth and incomplete pedigree information can lead to ambiguous allele calls, thereby increasing the likelihood of imputation errors (Chan et al., 2016). The OP simulation analysis is done in a way that is agnostic to the pollen parents identity of the OP genetic mapping population. In a real OP population, a major consideration for which paternal inference would be necessary for accurate QTL identification is if there are any genetic effects contributed by one or more pollen donors that influence the genetic mapping through additivity or epistasis or contribute significantly to the phenotypic variation (Dong et al., 2024). However, the simulations indicate that it would be feasible to accurately map QTL within an OP population if there are no additional QTL contributing to the trait of interest segregating from the pollen donors, particularly for large effect QTL.

Fine mapping a QTL in an OP population can be particularly challenging. Not only are the paternal genotypes unknown, but the genomic location of the QTL may also be entirely undefined prior to mapping, complicating efforts to identify the precise region of interest. If a QTL falls within a region of high LD or low recombination the ability to fine map in an F1 population could require restrictively large population sizes (Flint-Garcia et al., 2003). However, fine mapping is done as a way to further explore genetic regions and identify potential candidate genes and variants for QTL that have already been mapped (Fahrentrapp et al., 2013; Soriano et al., 2014) and these considerations should be taken into account when selecting the maternal parent for OP genetic mapping. A two-stage op mapping framework could be utilized in which a smaller population is first used to identify the QTL region, followed by mapping with additional F1s based on the local recombination rate in the identified region. Genetic mapping populations of OP F1 apple could be used as a way to more efficiently generate fine mapping populations after a large potential region of a QTL has been identified. In the case where the LD in the QTL region is high, using an OP F1 genetic mapping population may not be feasible, however, with lower LD regions or regions with higher recombination would allow fine mapping in an OP F1 population.

## 5 CONCLUSION

GWAS can be a useful QTL mapping method for apples that leverages the large amount of diversity found in core collections and germplasm repositories. However, the rapid LD decay, population genetic structure, and high heterozygosity in apples have restricted the widespread use of GWAS for QTL mapping in apples. Genetic mapping based on OP populations can accelerate the mapping of QTLs and development of reliable markers for marker assisted selection (MAS) for woody perennial crops. Markers tightly linked to a QTL are especially important for reducing linkage drag and to avoid the risk of losing the desired trait due to recombination in MAS (Hospital, 2009). This underscores the necessity of precise, fine-scale mapping to streamline the introgression of beneficial alleles while minimizing the inheritance of unwanted traits particularly when using wild species. The ability to use OP populations for QTL mapping as an alternative to traditional QTL mapping approaches has the potential to significantly reduce the labor and costs associated with growing and maintaining bi-parental populations. The enhanced genetic diversity in the OP populations will also facilitate the detection of QTLs that are effective across a much wider range of genetic backgrounds. Although, the simulations in this study were conducted solely in apples, further studies could extend the benefits of OP mapping to other perennial fruit tree crops.

GWAS has been valuable for QTL discovery in apples by leveraging the genetic diversity present in germplasm repositories and core collections. However, challenges such as rapid LD decay, high levels of population structure, and high heterozygosity limit its broader application in QTL mapping. In contrast, OP populations offer a practical alternative for genetic mapping in woody perennials. By utilizing natural pollination within genetically diverse orchards, OP mapping reduces the time, labor, and costs associated with maintaining large bi-parental populations.

Importantly, OP based mapping can accelerate the identification of tightly linked markers for MAS, which are critical for minimizing linkage drag and preserving desired traits when developing new varieties especially when incorporating alleles from wild relatives (Hospital, 2009). The increased genetic diversity captured in OP populations may also improve the detection of QTLs effective across diverse genetic backgrounds. While this study focused on apples, the mapping framework developed here has broad potential. Future research could expand its application to other perennial fruit tree crops, offering a more efficient path to trait discovery and marker development in complex, heterozygous species.

## DATA AVAILABILITY

The genotype by sequencing SNP data set for the populations GMAL4590, GMAL4591, GMAL4592, GMAL4593, GMAL4594, GMAL4595, and GMAL4596 used to calculate the genomic relationship matrix, simulate phenotypes, and perform marker-trait associations are made available on the Dryad database (https://doi.org/10.5061/dryad.15dv41p3c). The link to simulation script is available on GitHub at the following link: https://github.com/dh564/OP_Sim.

## ACKNOWLEDGEMENTS

This research was supported by the USDA-AFRI Plant Breeding for Agricultural Production (A1141) grant # 2023-67013-39303, the USDA-NIFA Specialty Crop Research Initiative (SCRI) grant 1023572 (subaward RC111414A) and the New York State Department of Agriculture & Markets, Apple Research & Development Pro-gram (ARDP), and the National Science Foundation National Research Traineeship (NRT) program under grant 1922551.

## DECLARATIONS

Not applicable

## AUTHOR CONTRIBUTIONS

A.K. conceptualized the research question. A.K. and K.R designed and supervised the research. D.H. performed all analysis and drafted the manuscript. D.H. A.K. and K.R revised, and finalized the manuscript. All authors read and approved the final version of the manuscript.

## Abbreviations

CIM: composite interval mapping
FDR: false discovery rate
GBS: genotyping-by-sequencing
GAPIT: Genome Association and Prediction Integrated Tool
GRM: genomic relationship matrix
GWAS: genome wide association studies
K-W: Kruskal-Wallis
LD: linkage disequilibrium
LG: linkage group
MLM: mixed linear model
MLMM: multi-locus mixed model
OP: open-pollinated
PGRU: Plant Genetic Resources Unit
QTL: quantitative trait locus.

