## Supplemental Tables (1-3) for "Development of an open-pollinated genetic mapping framework to facilitate the identification of QTL in apples"

**SUPPLEMENTAL DATA**

**Supplemental Table 1** Distance to the causative marker in base pairs. The average distance to the causative marker is reported across the 40 repeated simulations at each population size.

| **Fine Mapping Resolution (bp) on LG 1** | | | **Fine Mapping Resolution (bp) on LG 7** | | |
| --- | --- | --- | --- | --- | --- |
| n | $V_{G}$ = 0.5 | $V_{G}$ = 0.8 | n | $V_{G}$ = 0.5 | $V_{G}$ = 0.8 |
| 45 | 4.08E+06 | 4.65E+06 | 45 | 1.51E+07 | 8.28E+06 |
| 91 | 3.02E+06 | 8.92E+05 | 91 | 1.72E+06 | 1.30E+06 |
| 137 | 3.30E+06 | 1.53E+06 | 137 | 4.54E+06 | 1.06E+06 |
| 183 | 2.49E+06 | 1.29E+06 | 183 | 1.91E+06 | 4.35E+05 |
| 229 | 8.13E+05 | 7.01E+05 | 229 | 1.55E+06 | 2.54E+05 |
| 274 | 9.55E+05 | 9.35E+05 | 274 | 1.07E+06 | 4.92E+05 |
| 320 | 4.29E+05 | 2.44E+05 | 320 | 1.58E+06 | 5.03E+05 |
| 366 | 3.30E+05 | 4.07E+05 | 366 | 4.10E+05 | 2.05E+05 |
| 412 | 4.68E+05 | 1.13E+05 | 412 | 9.21E+05 | 9.44E+05 |
| 458 | 2.26E+05 | 4.80E+05 | 458 | 8.85E+05 | 6.61E+05 |
| 503 | 5.10E+05 | 5.92E+05 | 503 | 2.53E+05 | 1.98E+05 |
| 549 | 3.05E+05 | 5.59E+04 | 549 | 7.02E+05 | 8.40E+05 |
| 595 | 9.07E+05 | 3.93E+05 | 595 | 3.49E+05 | 3.39E+05 |
| 641 | 1.61E+06 | 5.04E+05 | 641 | 2.11E+05 | 4.18E+05 |
| 687 | 7.49E+05 | 1.75E+05 | 687 | 1.92E+05 | 2.80E+05 |

**Supplemental Table 2** False discovery rate (FDR) at each population size based on the 40 repeated simulations across the 15 different population sizes. Top of FormBottom of Form

| **FDR on LG1** | | | **FDR on LG7** | | |
| --- | --- | --- | --- | --- | --- |
| n | $V_{G}$= 0.5 | $V_{G}$= 0.8 | n | $V_{G}$= 0.5 | $V_{G}$ = 0.8 |
| 45 | 0.50 | 0.47 | 45 | 0.86 | 0.53 |
| 91 | 0.50 | 0.13 | 91 | 0.13 | 0.12 |
| 137 | 0.37 | 0.22 | 137 | 0.11 | 0.05 |
| 183 | 0.23 | 0.17 | 183 | 0.05 | 0.02 |
| 229 | 0.10 | 0.05 | 229 | 0.08 | 0.05 |
| 274 | 0.15 | 0.12 | 274 | 0.12 | 0.05 |
| 320 | 0.08 | 0.00 | 320 | 0.07 | 0.02 |
| 366 | 0.00 | 0.05 | 366 | 0.02 | 0.00 |
| 412 | 0.02 | 0.00 | 412 | 0.05 | 0.05 |
| 458 | 0.00 | 0.07 | 458 | 0.07 | 0.05 |
| 503 | 0.05 | 0.05 | 503 | 0.00 | 0.00 |
| 549 | 0.05 | 0.00 | 549 | 0.07 | 0.07 |
| 595 | 0.02 | 0.07 | 595 | 0.02 | 0.02 |
| 641 | 0.14 | 0.02 | 641 | 0.00 | 0.02 |
| 687 | 0.07 | 0.05 | 687 | 0.00 | 0.02 |

**Supplemental Table 3** Average power to detect the simulated QTL across the 40 repeated simulations at each population size.

| **Average Power LG1** | | | **Average Power LG7** | | |
| --- | --- | --- | --- | --- | --- |
| n | $V_{G}$= 0.5 | $V_{G}$= 0.8 | n | $V_{G}$= 0.5 | $V_{G}$ = 0.8 |
| 45 | 0.00 | 0.10 | 45 | 0.08 | 0.30 |
| 91 | 0.08 | 0.50 | 91 | 0.35 | 0.73 |
| 137 | 0.45 | 0.68 | 137 | 0.33 | 0.88 |
| 183 | 0.35 | 0.83 | 183 | 0.68 | 0.93 |
| 229 | 0.75 | 0.88 | 229 | 0.78 | 0.95 |
| 274 | 0.65 | 0.73 | 274 | 0.85 | 0.98 |
| 320 | 0.83 | 0.88 | 320 | 0.88 | 1.00 |
| 366 | 0.83 | 0.90 | 366 | 0.93 | 0.98 |
| 412 | 0.80 | 0.93 | 412 | 0.88 | 1.00 |
| 458 | 0.88 | 0.93 | 458 | 0.95 | 1.00 |
| 503 | 0.85 | 1.00 | 503 | 1.00 | 1.00 |
| 549 | 0.93 | 0.98 | 549 | 0.98 | 1.00 |
| 595 | 0.98 | 0.95 | 595 | 1.00 | 1.00 |
| 641 | 0.90 | 0.98 | 641 | 0.98 | 1.00 |
| 687 | 0.95 | 0.95 | 687 | 1.00 | 1.00 |
